# Identification of biomarkers and pathways of mitochondria in sepsis patients

**DOI:** 10.1101/2021.03.29.437586

**Authors:** Gongsheng Yuan

**Affiliations:** Department of Physiology and Pathophysiology, School of Basic Medical Sciences, Fudan University, Shanghai, China

## Abstract

Sepsis is a life-threatening condition associate with significant morbidity and mortality, but limited treatment. Mitochondria are recently recognized to be related to the pathophysiology of sepsis, and mitochondria could serve as a potential drug target. In our study, we aim to identify biological functions and pathways of mitochondria during the processes of sepsis by using a bioinformatics method to elucidate their potential pathogenesis. The gene expression profiles of the GSE167914 dataset were originally created by using the Nanostring nCounter Elements™ TagSet preselected for mitochondrial biogenesis and function panel. The biological pathways were analyzed by the Kyoto Encyclopedia of Genes and Genomes pathway (KEGG), Gene Ontology (GO), and Reactome enrichment. KEGG and GO results showed the Neurodegeneration pathways such as Huntington and Parkinson pathways were mostly affected in the development of sepsis. Moreover, we identified several mitochondrial genes including TOMM40, TOMM20, TIMM22, TIMM10, TIMM17A, TIMM9, TIMM44 were involved in the regulation of protein translocation into mitochondria. Further, we predicted several regulators that had the ability to affect the mitochondria during sepsis by L1000fwd analysis. Thus, this study provides further insights into the mechanism of mitochondrial function during sepsis.

## Introduction

Sepsis is caused by the dysregulated host response to infections from tissue damage or organ dysfunction^1^. Sepsis is a serious disease with a mortality of 15-20%, which is characterized not only by the upregulation of inflammation but also by the strong immune suppression^2^. Understanding the pathophysiology of sepsis may aid in the development of novel therapies^3^. The effects of impaired cellular functions such as mitochondrial dysfunction and cell death mechanism in sepsis-associated organ dysfunction are beginning to be unrevealed^4^.

The significance of mitochondrial dysfunction in sepsis is previously recognized^5^. As the mitochondrion is a critical organelle for multiple cellular processes such as adenosine triphosphate (ATP) production, intracellular calcium homeostasis, and the production of ROS and some hormones^6, 7^. Moreover, mitochondria are also associated with triggering the intrinsic pathway of apoptosis, which is caused by the outer membrane permeabilization^8^. The mitochondrial functions are changed during sepsis, which includes reduced oxidative phosphorylation, increased ROS production, and altered mitochondrial biogenesis^9^. One hypothesis to explain the dysregulated mitochondrial function is that the decreased oxidative phosphorylation might contribute to the reduced production of potentially harmful ROS^10^. Mitophagy and mitochondrial biogenesis might be involved in sepsis as a mechanism to reduce the harmful effects of mitochondrial dysfunction^5^. Therefore, during the progression of sepsis, there are various mitochondrial alterations at different times. However, what kind of alteration affects the mitochondrial functions is unclear. Thus, studying the regulation of mitochondria and the related pathways may be a convincing strategy to study the mechanism of sepsis. In this study, we investigated the effect of mitochondrial alterations in sepsis patients. We identified several DEGs, candidate inhibitors, and the relevant biological process in sepsis patients by utilizing comprehensive bioinformatics analyses. The functional enrichment analysis and protein-protein interaction analysis were used for discovering significant gene nodes. These key genes and signaling pathways may be essential to therapeutic interventions of sepsis.

## Methods

### Data resources

The GSE167914 dataset was obtained from the GEO database (http://www.ncbi.nlm.nih.gov/geo/). The data was produced by Nanostring nCounter Elements™ TagSet preselected for mitochondrial biogenesis and function panel, Faculty of Medicine, Universitas Tarumanagara, Jakarta Barat, DKI Jakarta, Indonesia. RNA-Seq analysis was performed using peripheral blood of infection and sepsis patients as well as healthy controls.

### Data acquisition and preprocessing

The GSE167914 dataset that contains gene expression related to mitochondrial function from the peripheral blood of infection and sepsis patients as well as healthy controls was conducted by R script^11, 12^ We used a classical t test to identify DEGs with P<.01 and fold change ≥1.5 as being statistically significant.

### Gene functional analysis

Gene Ontology (GO) is a community-based bioinformatics resource that contains the model Biological Process, Molecular Function, and Cellular Component. Kyoto Encyclopedia of Genes and Genomes (KEGG) database is a useful tool that integrates functional information, biological pathways, and sequence similarity^13^. GO and KEGG pathway analyses were performed by utilizing the Database for Annotation, Visualization, and Integrated Discovery (DAVID) (http://david.ncifcrf.gov/) and Reactome (https://reactome.org/). P<0.05 and gene counts >10 were considered statistically significant.

### Module analysis

Molecular Complex Detection (MCODE) of Cytoscape software was used to study the connected regions in protein-protein interaction (PPI) networks. The significant modules and clusters were selected from the constructed PPI network using MCODE and String (https://string-db.org/). The pathway enrichment analyses were performed by using Reactome, and P<0.05 was used as the cutoff criterion.

### Reactome analysis

The Reactom pathway (https://reactome.org/) was used to obtain the visualization, interpretation, and analysis of potential pathways. P<.05 was considered statistically significant.

## Results

### Identification of DEGs of mitochondria from the blood of sepsis patients

The peripheral blood of sepsis patients and healthy controls were harvested to analyze the DEGs (differentially expressed genes) of mitochondria. Patients fulfilling infection or sepsis criteria were recruited from the emergency department (Faculty of Medicine, Universitas Tarumanagara, Velma Herwanto, Indonesia). A total of 56 genes were identified to be differentially expressed in sepsis patients with the threshold of P<0.05. The top 10 up- and down-regulated genes are listed in table 1.

**Table 1.**

| **Entrez gene** | **Gene Symble** | **Fold-change** | **Regulation** |
| --- | --- | --- | --- |
| Top 10 down-regulated DEGs |  |  |  |
| 706 | TSPO | -1.45828702346694 | Down |
| 26521 | TIMM8B | -1.43210769844688 | Down |
| 51100 | SH3GLB1 | -1.28619452774405 | Down |
| 29957 | SLC25A24 | -1.01302822644703 | Down |
| 55288 | RHOT1 | -1.01271887616278 | Down |
| 788 | SLC25A20 | -0.901108073452252 | Down |
| 84134 | TOMM40L | -0.857069247581503 | Down |
| 10245 | TIMM17B | -0.829410945662233 | Down |
| 3939 | LDHA | -0.800959299777023 | Down |
| 67126 | ATP5E | -0.784378992381854 | Down |
| Top 10 up-regulated DEGs |  |  |  |
| 7157 | TP53 | 1.12448582743376 | up |
| 23082 | PPRC1 | 1.03266502881293 | up |
| 596 | BCL2 | 0.994531085428525 | up |
| 66043 | ATP5D | 0.949586740919676 | up |
| 7351 | UCP2 | 0.864981370741031 | up |
| 84172 | POLR1B | 0.728977834365151 | up |
| 3418 | IDH2 | 0.644515388301858 | up |
| 10128 | LRPPRC | 0.642054261768502 | up |
| 23 | ABCF1 | 0.632440787164533 | up |
| 10953 | TOMM34 | 0.623887917139965 | up |

### KEGG analysis of DEGs of mitochondria from the blood of sepsis patients

To identify the biological functions and potential mechanisms of DEGs of mitochondria from sepsis patients and healthy controls, we performed KEGG pathway enrichment analysis and created a visual graph (Supplemental Table S1). KEGG pathway (http://www.genome.jp/kegg/) is a gene collection for exploring the molecular interaction, reaction, and relation networks. Our study showed top ten enriched KEGG pathways including “Neurodegeneration”, “Platinum drug resistance”, “p53 signaling pathway”, “Huntington disease”, “Mitophagy”, “Apoptosis”, “Parkinson disease”, “Apoptosis-multiple species”, “Measles”, and “Colorectal cancer” (Figure 1).

**Figure 1.**
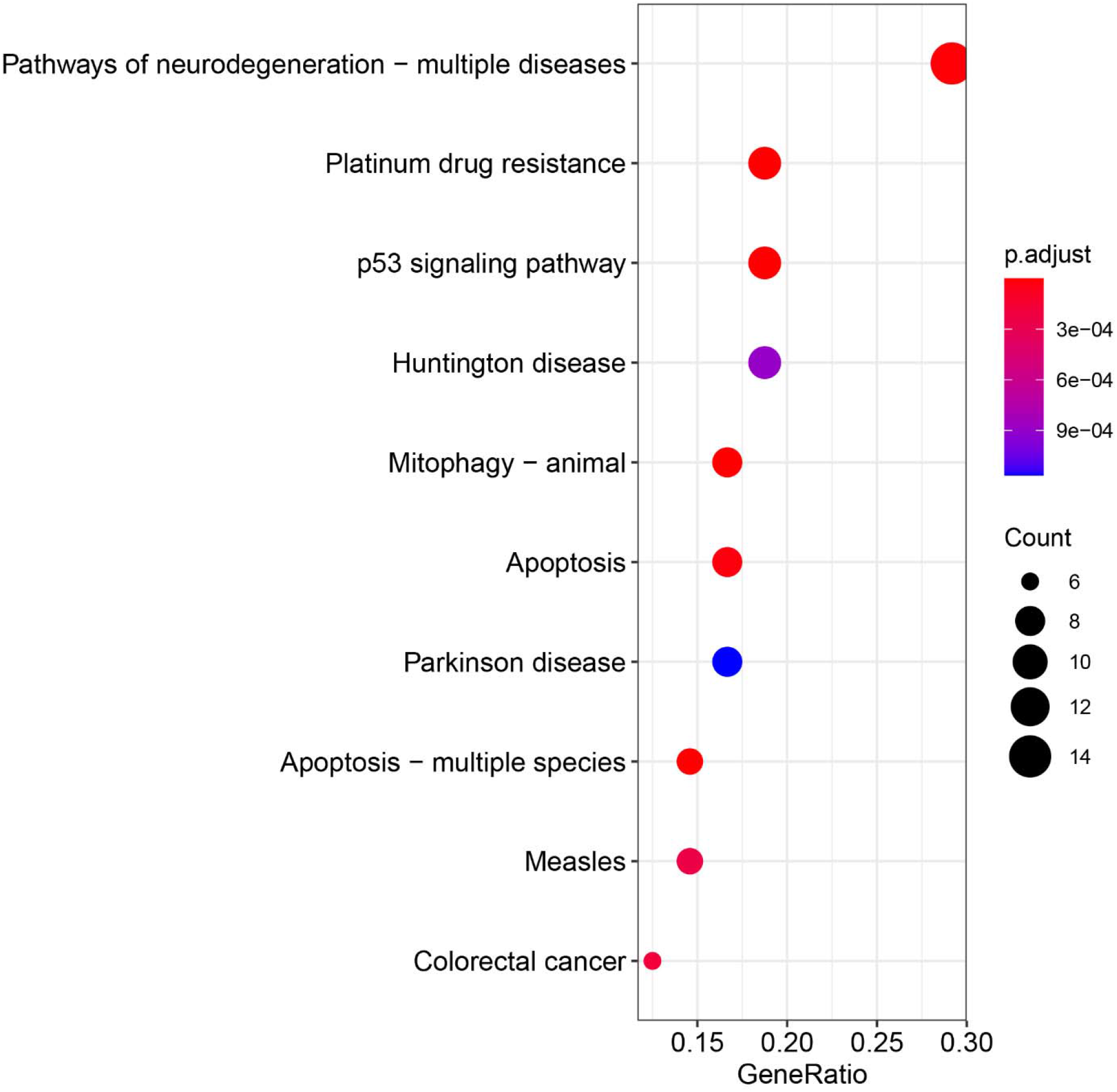
The KEGG pathways enriched by the DEGs. DEGs =differentially expressed genes, KEGG = Kyoto Encyclopedia of Genes and Genomes.

### GO analysis of DEGs from the blood of sepsis patients

Gene Ontology (GO) analysis is a commonly used tool for classifying genes, which includes cellular components (CC), molecular functions (MF), and biological processes (BP). Here, we identified the top ten cellular components including “mitochondrial inner membrane”, “organelle outer membrane”, “outer membrane”, “mitochondrial outer membrane”, “mitochondrial protein complex”, “intrinsic component of organelle membrane”, “intrinsic component of mitochondrial membrane”, “integral component of organelle membrane”, “integral component of mitochondrial membrane”, and “intrinsic component of mitochondrial outer membrane” (Figure 1). We then identified the top ten biological processes: “mitochondrial transport”, “protein localization to mitochondrion”, “establishment of protein localization to mitochondrion”, “protein targeting to mitochondrion”, “mitochondrial transmembrane transport”, “mitochondrial membrane organization”, “ATP transport”, “adenine nucleotide transport”, “purine ribonucleotide transport”, and “purine nucleotide transport” (Figure 1). We also identified the top ten molecular functions: “organic anion transmembrane transporter activity”, “anion transmembrane transporter activity”, “ATP transmembrane transporter activity”, “adenine nucleotide transmembrane transporter activity”, “purine ribonucleotide transmembrane transporter activity”, “purine nucleotide transmembrane transporter activity”, “nucleotide transmembrane transporter activity”, “organophosphate ester transmembrane transporter activity”, “nucleobase-containing compound transmembrane transporter activity”, and “carbohydrate derivative transmembrane transporter activity” (Figure 2 and Supplemental Table S1).

**Figure 2.**
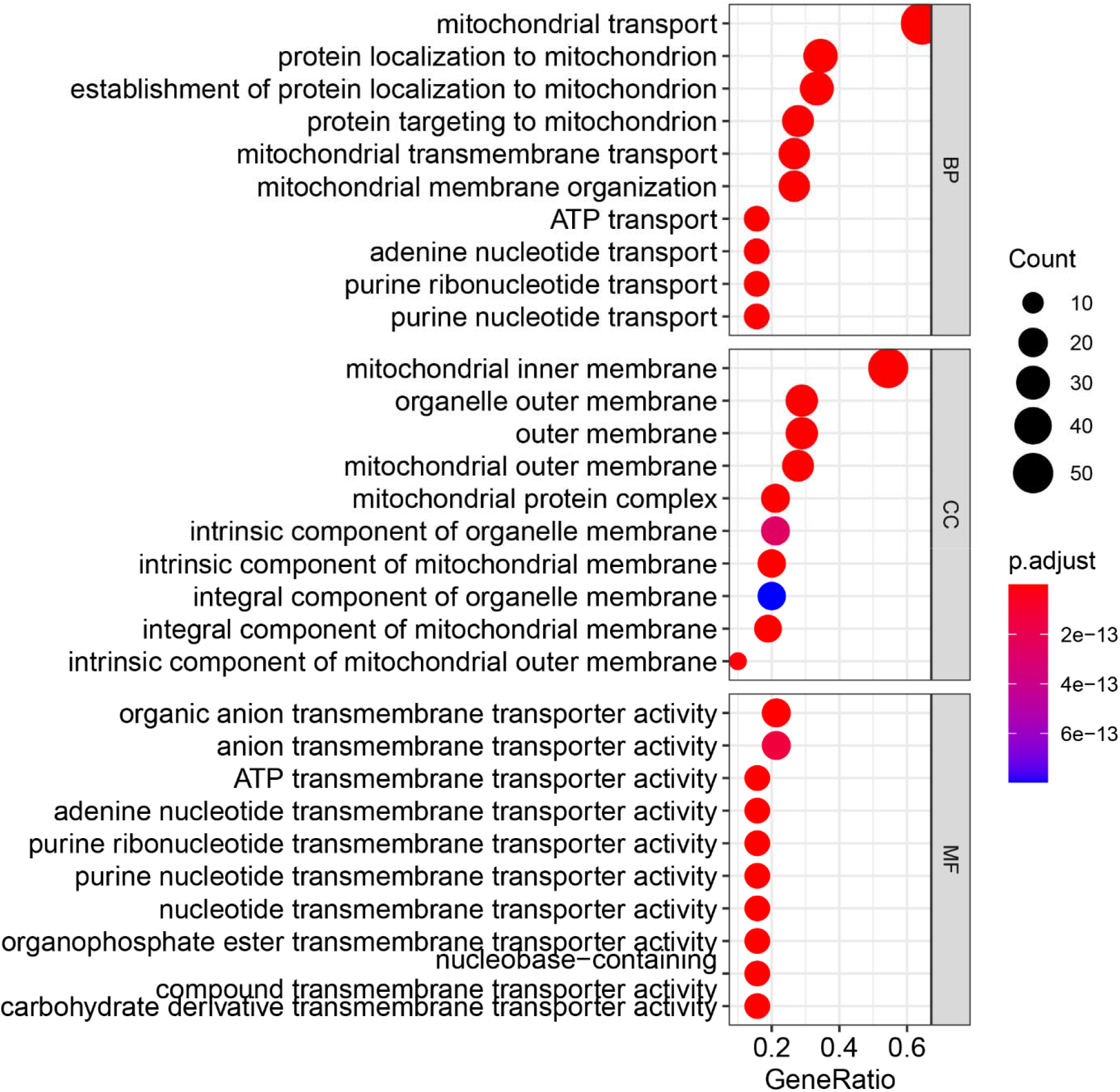
The biological process (BP), cellular component (CC), and molecular function (MF) terms enriched by the DEGs.

### PPI network and Module analysis

We constructed PPI networks to analyze the relationships of DGEs at the protein level. The criterion of combined score >0.7 was chosen and the PPI network was constructed by using 55 nodes and 106 interactions. Among these nodes, the top ten genes with the highest scores are shown in Table 2. The significant modules of DEGs of mitochondria from the blood of sepsis patients were selected to show the functional annotation (Figure 3).

**Figure 3.**
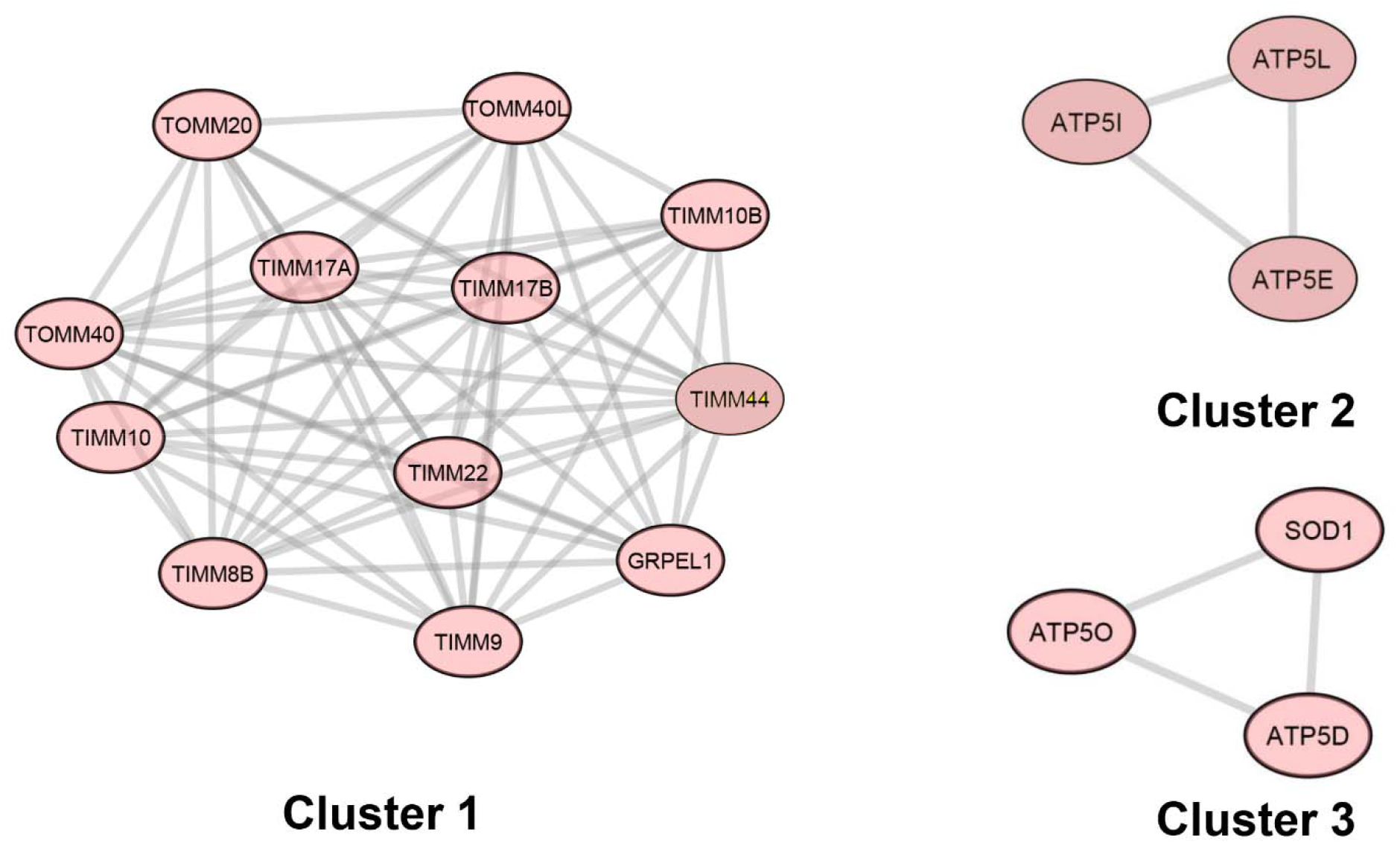
Top three modules (Cluster1-3) from the blood of sepsis patients

**Table 2.** Top ten genes demonstrated by connectivity degree in the PPI network

| Gene Symbol | Gene title | Degree |
| --- | --- | --- |
| TOMM40 | translocase of outer mitochondrial membrane 40 | 16 |
| TOMM20 | translocase of outer mitochondrial membrane 20 | 12 |
| TIMM22 | translocase of inner mitochondrial membrane 22 | 12 |
| TIMM10 | translocase of inner mitochondrial membrane 10 | 12 |
| TIMM17A | translocase of inner mitochondrial membrane 17A | 11 |
| TIMM17B | translocase of inner mitochondrial membrane 17B | 11 |
| CYCS | cytochrome c, somatic | 10 |
| TIMM9 | translocase of inner mitochondrial membrane 9 | 9 |
| TIMM44 | translocase of inner mitochondrial membrane 44 | 9 |
| TAZ | tafazzin | 7 |

### Reactome Pathway analysis

To further understand the potential functions of DEGs, we also identified several signaling pathways by using Reactome Pathway Database. The top ten signaling pathways include “Mitochondrial protein import”, “Protein localization”, “Mitochondrial biogenesis”, “Cristae formation”, “The citric acid (TCA) cycle and respiratory electron transport”, “Formation of ATP by chemiosmotic coupling”, “Organelle biogenesis and maintenance”, “Intrinsic Pathway for Apoptosis”, “Respiratory electron transport, ATP synthesis by chemiosmotic coupling, and heat production by uncoupling proteins”, and “Activation of PUMA and translocation to mitochondria” (Supplemental Table S2). We then constructed the visual reaction map according to the signaling pathways (Figure 4).

**Figure 4.**
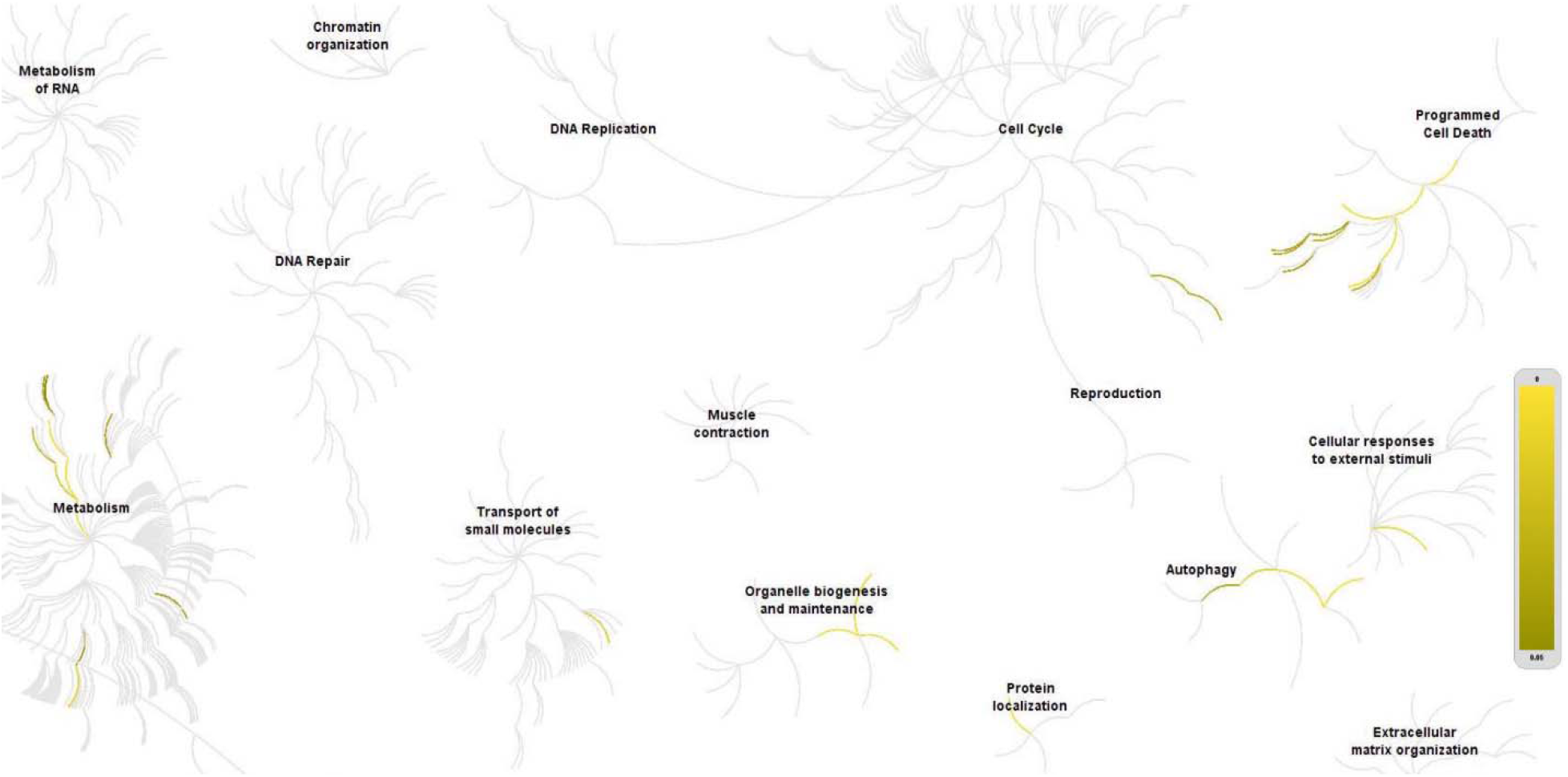
The Reactom pathway visualization map. Input genes are represented by the top significantly changed genes obtained from the GSE167914 dataset (P <0.01). The yellow color represents the most relevant signaling pathways.

### Potential inhibitors for the treatment of sepsis

To further know the potential inhibitors for the treatment of sepsis, we introduced the L1000 fireworks display system that can predict bioactive molecules. The system indicated the potential pathways that may be inhibited. We selected the top ten molecules according to the DEGs and the inhibitor map: “BRD-K39757396”, “BRD-K39829853”, “veliparib”, “BRD-K32862555”, “halcinonide”, “DG-041”, “HC-toxin”, “UK-356618”, “XMD-1150”, and “GSK-461364” (Figure 5 and Supplemental Table S3).

**Figure 5.**
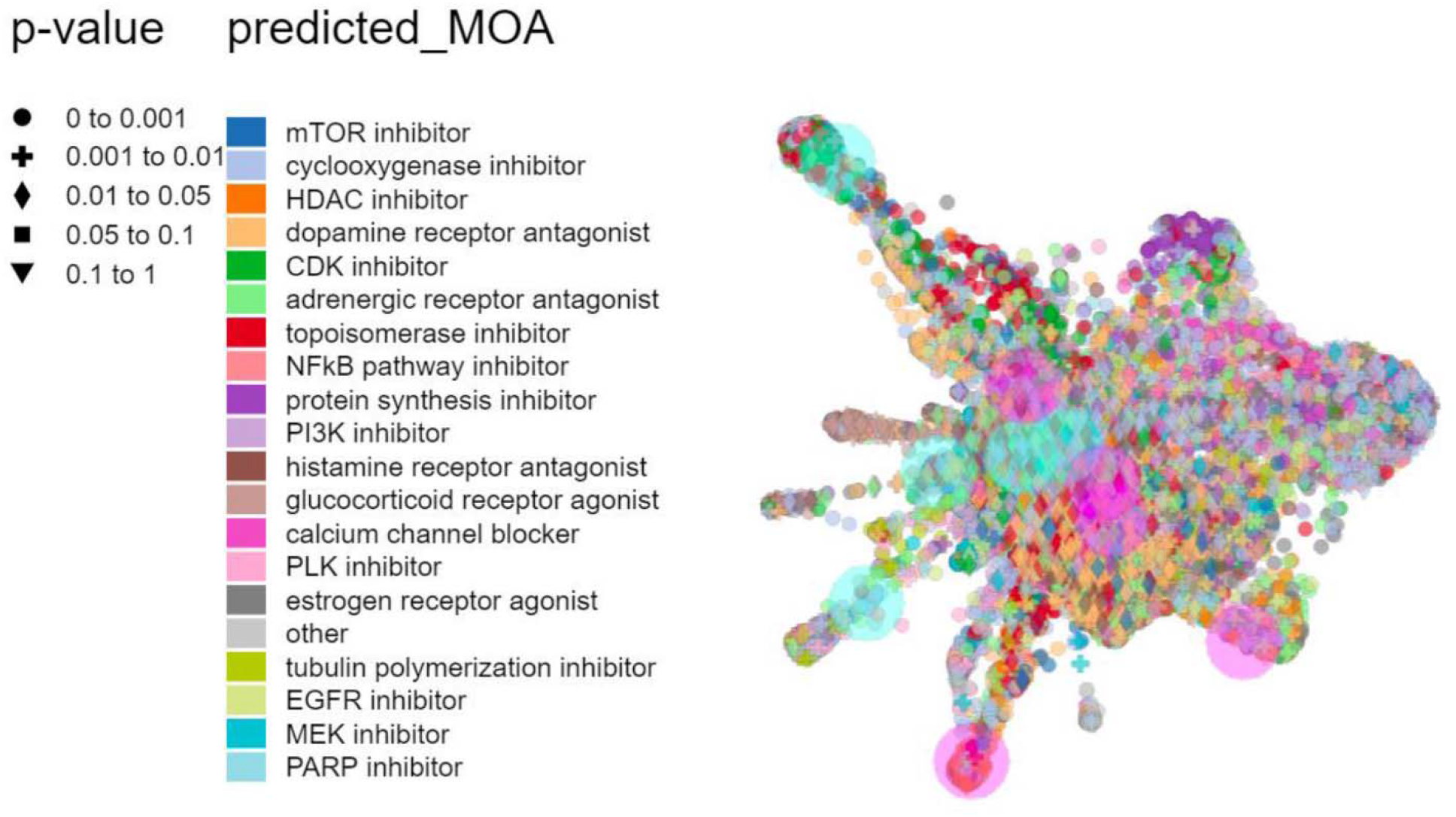
Inhibitors by L1000FDW visualization. Input genes are represented by the significantly changed genes obtained from the GSE167914 dataset. Dots are the Mode of Action (MOA) of the respective drug.

## Discussion

Sepsis is a kind of inflammation caused by a serious response of the immune system and finally led to multiorgan failure and death^14^. Sepsis is related to the activation of innate immunity through various pathological processes^15^. For example, the NF-κB signaling pathway as the inflammation center in various diseases^16, 17^ is activated by the pathogen-associated molecular patterns (PAMPs) during the beginning of sepsis^18^. Mitochondria provide energy to cells and produce ATP through the basic process of respiration. Numerous cells respond to stresses such as cytokines by triggering mitochondria-dependent signals that initiate cellular protective responses^19^. Thus, based on the mitochondrial study on sepsis patients, our study may provide gene evidence for clinical trials of sepsis.

To understand the effects of mitochondria in sepsis, we analyzed the mitochondrial respiration and gene expression related to mitochondrial functions from the peripheral blood of sepsis patients and healthy controls. Ten proteins were selected according to the PPI network analysis, which may be important during the development of sepsis. TOMM40 is associated with apolipoprotein E (APOE), which further influences age-related memory performance during Alzheimer’s disease^20^. TOMM20 promotes proliferation, resistance to apoptosis, and chemicals^21^. In melanoma cells, enhanced ROS leads to the oxidation and oligomerization of the mitochondrial outer membrane protein Tom^22^. TIM22 is the complex in the mitochondrial inner membrane^23^. Mitochondrial acylglycerol kinase assembles with TIMM22 to support the import of a subset of multi-spanning membrane proteins^24^. TIMM10 is identified as the blood-based biomarkers for pulmonary tuberculosis by modeling and mining molecular interaction networks^25^. TIM17A is the stress-regulated subunit of the translocase of TIM23, which is downregulated by the eukaryotic initiation factor 2α (eIF2α)^26^. TIM9 and TIM44 are responsible for the transport of proteins across the inner membrane^27^.

KEGG and GO analyses indicated that neurodegeneration was the main pathological process during sepsis. The KEGG analysis showed “Neurodegeneration”, “Huntington disease”, and “Parkinson disease” were the most phenotypes during sepsis, which suggested that the brain or nerve damage was the most affected part during the infection. Similarly, Catherine N Widmann reported that sepsis could cause an increase in the permeability of the blood-brain barrier, which may lead to a rapid decline in cognitive function or coma^28^. Thus, sepsis may increase the brain’s susceptibility to neurodegenerative disease. Moreover, we also found the cell death-related signaling processes “p53 signaling pathway”, “Mitophagy”, and “Apoptosis” were involved in sepsis. These signaling pathways were widely enhanced in the damaged cells or cancers^29–34^. Interestingly, these signaling pathways are regulated by the circadian genes such as Clock^16, 35^. Circadian clocks play important roles in the physiological and pathophysiological processes^36–40^, such as controlling the immune checkpoint pathways in immune cells^41^. The GO analysis also showed the mitochondrial activity was enhanced during sepsis. Processes like “mitochondrial transport”, “protein localization to mitochondrion”, “establishment of protein localization to mitochondrion” were associated with the mitochondrial function, which suggested that sepsis could affect the translocation function of mitochondria.

In summary, we identified the potential pathways in sepsis patients by analyzing the mitochondrial gene functions. Neurodegeneration diseases such as Huntington’s disease and Parkinson’s disease are the mainly triggered diseases during sepsis. Future studies will focus on the administration of potential mitochondrial regulators on clinical trials. This study thus provides further insights into the treatment of sepsis, which may facilitate drug development.

